# Transition from actin-driven to water-driven cell migration dependson external hydraulic resistance

**DOI:** 10.1101/275123

**Authors:** Y. Li, S.X. Sun

## Abstract

*In vivo*, cells can reside in diverse physical and biochemical environments. For example, epithelial cells typically live in a two-dimensional (2D) environment while metastatic cancer cells can move through dense three-dimensional (3D) matrices. These distinct environments impose different kinds of mechanical forces on cells, and thus potentially can influence the mechanism of cell migration. For example, cell movement on 2D flat surfaces is mostly driven by forces from focal adhesion and actin polymerization, while in confined geometries, it can be driven by water permeation. In this work, we utilize a two-phase model of the cellular cytoplasm, where the mechanics of the cytosol and the F-actin network are treated on an equal footing. Using conservation laws and simple force balance considerations, we are able to describe the contribution of water flux, actin polymerization and flow, and focal adhesions to cell migration in both 2D surfaces and in confined spaces. The theory shows how cell migration can seamlessly transition from a focal adhesion- and actin-based mechanism on 2D surfaces to a water-based mechanism in confined geometries.

## 1 Introcudtion

Animal cell migration is a complex process orchestrated by actin dynamics, focal adhesions, and also water flux [1, 2]. However, how these elements are added together to obtain the observed cell speed is less clear. For example, cell migration on two-dimensional (2D) surfaces is mostly driven by forces from actin polymerization and focal adhesions [3] and there has been extensive work on modeling actin-driven cell migration on 2D surfaces [4, 5, 6], while cells in confined geometries can be driven by water permeation [7]. Moreover, cells in confined channels show a diversity of behavior: some cells such as MDA cells show reduced migration speed when actin is disrupted [8, 9], while for others such as S180, the migration speed is unaffected [7]. In addition to varying responses to actin inhibition, cell movement in confinement appears to be sensitive to the hydraulic resistance [10]. Even more complex are cells in three-dimensional (3D) collagen matrices where they develop long protrusions that interact with the collagen fibers, pores, and interstitial fluid [11, 12, 13, 14, 15]. Depending on the cell shape, the nucleus may also play a significant role in propelling the cell [16]. Thus, there are diverse mechanisms driving cell migration [17]. We would like to understand whether there are unified physical principles and mechanisms giving rise to the wide range of observed cell behavior. Can we explain the impact of the physical environment on the speed of cell migration? In this paper, by focusing on the combined contributions from actin dynamics and water flow, we develop a general model to understand mechanisms of cell migration in 3D, 2D, and one-dimensional (1D) environments (Fig. 1A-C). In all these cases, we examine an effective one-dimensional volume element of a cell and compute the leading edge cell speed for different actin and water flow dynamics. We find that the hydraulic environment of the cell has a counterintuitive impact on the cell speed and determine the contribution of actin and water to the observed cell movement. In particular, cells can speedup if the coefficient of external hydraulic resistance increases, even when there is no change in the molecular elements driving migration. These results explain the diversity of observed cell migration mechanisms, and suggest that cells moving in 3D matrices are not only influenced by collagen fibers, but also the hydrodynamic environment.

**Figure 1:**
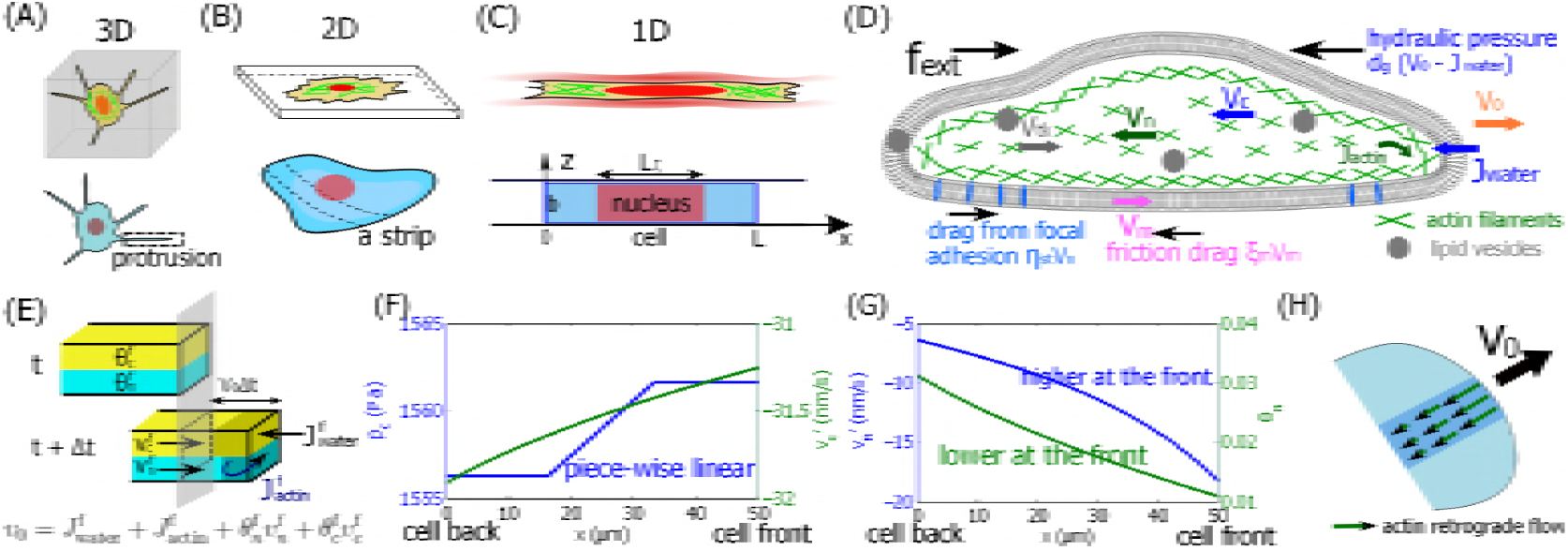
A two-fluid-phase cell migration model. (A) Schematics of a cell in a 3D collagen gel. The thin protrusion can be regarded as a 1D structure. (B) Schematics of a cell on a 2D substrate. A strip of a cell on a 2D substrate can be regarded a 1D structure. (C) Schematics of a cell in a confined space. In this case, the cell is essentially 1D. The model represents a cell (or a strip) with length *L* and width *b*. (D) Diagram of the relevant forces that contribute to cell migration. *v*_*n,c*_ is the F-actin and cytosol phase velocity, respectively. *v*_t*k*_ is the possible vesicle trafficking rate from the back to the front. The actin phase form focal adhesion with the environment, resulting in frictional drag force *η*_st_*v*_*n*_. Membrane movement also generates frictional force *ξ*_*m*_*v*_*m*_. As the cell displaces external water, the hydraulic resistance can be expressed as *d*_*g*_(*v*_0_ *- J*_water_), where *v*_0_ is the cell boundary velocity. (E) A unit cross-sectional area of the cell leading edge. An exact kinematic relation for two-phase cell migration from time *t* to *t* + Δ*t* can be derived from mass balance. Each phase is attached to the cell leading edge (no void space) so that the cell velocity is related to the velocity of each phase. 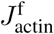 is the actin polymerization rate at the leading edge. 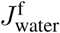 is the water influx rate at the leading edge. *θ*_*c,n*_ is the volume fraction of the cytosol (actin) phase, respectively. (F) Computed intracellular hydrostatic pressure field (left axis) and cytosol velocity field with respect to the cell frame (right axis). 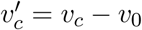. (G) Computed intracellular actin velocity field with respect to the cell frame (left axis) and its volume fraction (right axis). 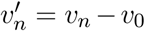. (H) Diagram of the actin retrograde flow predicted by the model.

## 2 A two-phase model of cell migration

Here we discuss briefly the two phase model of cell migration; more details can be found in the supplemen-tary material (SM). In our model we treat the cytosol (*c*, which is essentially water) and the actin network (*n*) as two fluid phases interacting with each other [18] and with the environment. The essential elements in the model are illustrated in Fig. 1D. The volume fraction and velocity of the cytosol (actin network) phase are *θ*_*c*_(*θ*_*n*_) and *v*_*c*_(*v*_*n*_), respectively. If we regard the actin network as isotropic, then it also has an effective pressure that is different from the cytosol phase due to active contractions from myosin (SM). The actin network is linked to the underlying substrate through focal adhesions and integrins [19]. As the actin network flows, this linkage transmits a force from the environment to the actin network and thus onto the entire cell. For low actin flow velocities, the force from focal adhesions is approximately proportional to the velocity of actin flow [20], i.e., *f*_st_ = *-η*_st_*v*_*n*_, where *η*_st_ is the coefficient of focal adhesion friction. Factors that can influence *η*_st_ include the substrate stiffness [21, 22] and the size [23] and density [24] of adhesions.

For moving cells, new actin filaments are polymerized at the cell boundary with a rate *J*_actin_. Similarly water influx, *J*_water_, also occurs at the cell boundary. In this model *J*_actin_ is a parameter, and *J*_water_ is calculated from the chemical potential difference of water, i.e., *J*_water_ = *-α*(Δ*P -* ΔΠ), where Δ*P* and ΔΠ are, respectively, the hydrostatic and osmotic pressure differences across the cell membrane, and *α* is a coefficient of membrane water permeability. The cell can control ΔΠ by generating ion fluxes at the cell boundary, and therefore we take it as a parameter. Δ*P* must be computed from mechanical considerations.

As the cell boundary moves at velocity *v*_0_, the cell may experience an extracellular hydraulic resistance (in units of pressure because this force is exerted on an area element). When *v*_0_ /= *J*_water_, the cell boundary must push the external fluid at velocity *v*_*\**_ = *v*_0_ *- J*_water_. In the extreme case of a cell in a confined channel, the cell must push the complete column of water in front of the cell at the velocity *v*_*\**_, and the friction between the extracellular fluid and the channel wall generates a large hydraulic resistance. This hydraulic resistance can be expressed as a linear function of *v*_*\**_, i.e., *P*_*\**_ = *d*_*g*_(*v*_0_ *- J*_water_), where *d*_*g*_ is a coefficient to be determined by the extracellular geometry and flow properties. In addition, the cell may experience external forces from extracellular objects such as mounted cantilever [25]. We use *f*_ext_ to account for this effective force per unit area (again in units of pressure).

To avoid complications associated with cell geometry, in this paper we mainly discuss a 1D volume element of a moving cell. For example, for cells in 3D collagen matrices (Fig. 1A), the thin protrusions can be regarded as 1D structures. For cells on 2D substrates (Fig. 1B), we model a 1D strip of the cell. In this case, we only consider velocities and forces perpendicular to the cell leading edge. For cells in a confined space (Fig. 1C), the actin network and water flows can be directly modeled in 1D. In this case, the cell nucleus provides additional drag forces on the cytoplasm as the flows pass around or through (for the water phase) the nucleus (SM).

Within the 1D framework, the cell boundary is reduced to a front (‘f’) and a back (‘b’). Since there are no void spaces, the cell boundary velocity is closely related to the boundary velocity of each phase. At the leading edge of the cell, as illustrated in Fig. 1E, boundary kinematic relations read as 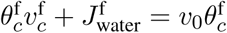 and 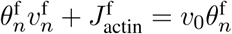. Therefore the velocity of the cell boundary is

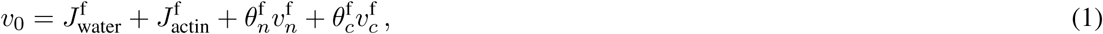

where we have used *θ*_*c*_ +*θ*_*n*_ = 1. Eq. 1 suggests that both water flux and actin polymerization can potentially contribute to cell migration [26]. In order to predict *v*_0_ we must know 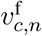, which, as we will show, can vary depending on the physical environment of the cell. The flow velocities, *v*_*c,n*_(*x*), can be computed from force balances of each phase.

During cell migration, the cell membrane has a translational velocity, *v*_*m*_, with respect to the surroundings. This motion leads to a frictional force between the cell and the surroundings. This force can be expressed as *f*_*m*_ = *-ξ_m_v_m_*, where *ξ*_*m*_ is the coefficient of frictional drag. *v*_*m*_ and *v*_0_ are typically equivalent, but they can also be different if we include the velocity of membrane growth or extension from vesicle trafficking [27], *v*_tk_. In this case, the apparent velocity of the cell boundary, *v*_0_, is formally a sum of the translational velocity of the lipid membrane and the velocity from vesicle trafficking, i.e., *v*_0_ = *v*_*m*_ + *v*_tk_. *v*_tk_ depends on the cell membrane tension, *τ* [28, 29]. Note that vesicle trafficking does not affect Eq. 1, which is only based on the assumption that the actin network phase is always attached to the membrane. This assumption holds during typical tissue cell migration. One violation of such condition is blebbing motility [30], where the membrane extends without actin. While blebbing motility can be analyzed within the two-phase framework, the details are more complicated. We will not discuss this case here.

The cell boundary velocity *v*_0_ is obtained from the entire coupled system where fluid pressure, velocities, volume fractions, boundaries conditions, and force balances are all considered. An analytic approximation for *v*_0_ when *v*_*m*_ = *v*_0_ is (see SM for more information)

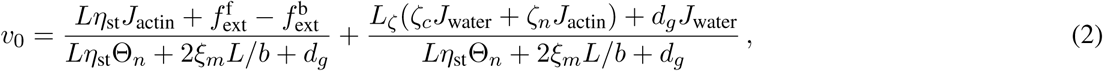

where *L* and *b* are the length and width of the cell (or a strip of cell), respectively. *L*_*ζ*_ is the length of the nucleus (Fig. 1C). *ζ*_*c*_ and *ζ*_*n*_ are, respectively, the effective coefficients of drags on the cytosol and actin network from the nucleus. Θ*_n_* is the average volume fraction of the actin phase satisfying 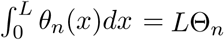. At steady-state with constant cell volume, 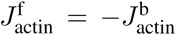 and 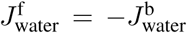; we thus use the notations 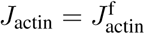 and 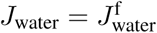 to represent the magnitude of these fluxes in the cell.

## 3 Results

### 3.1 Nucleus can increase the intracellular pressure difference

Fig. 1F-G shows a typical distribution of the intracellular hydrostatic pressure, volume fraction of the actin phase, and velocities of the two phases for a migrating cell with respect to the cell moving frame (see SM for parameters). The velocity of the cell edge and the membrane translation are predicted the same, *v*_0_ = *v*_*m*_ = 12 nm/s. Therefore, in this case vesicle trafficking has a negligible effect. This is because the membrane tension difference between the cell front and back is negligible (see SM and a later section). For the assumed osmotic pressure gradient, the water flux into the cell at the front is *J*_water_ = 31 nm/s while the actual cell velocity is much smaller. Since *v*_0 /=_ *J*_water_, the cell displaces the extracellular fluid as it migrates.

The average intracellular hydrostatic pressure, *P*_*c*_, is determined by the average osmolarity difference across the cell. In confined channels, the nucleus also influences *P*_*c*_, giving rise to a piece-wise linear pressure profile. Larger effective drag from the nucleus on the cytosol, i.e., larger *ζ*_*c*_, induces higher pressure difference between the front and back of the cell. When *ζ*_*c*_ = 0, then *P*_*c*_ is a smooth function over the entire cell length and the difference of *P*_*c*_ across the cell is reduced (see Fig. S2). This result is consistent with the ‘nuclear piston’ mode of migration where the pressure gradient of the nucleus can influence cell speed [16]. The relative velocity of the cytosol, 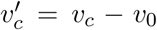 shown in Fig. 1F, is determined mostly by the boundary condition, i.e., 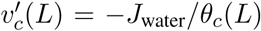. Higher 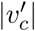 at the back of the cell is consistent with the lower cytosolic volume fraction since the product 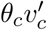 must be a constant from the mass conservation.

### 3.2 Retrograde flow is prominent at the cell leading edge

The relative velocity of the actin network, 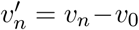 shown in Fig. 1G, is largely determined by the rate of actin (de)polymerization at the boundaries, i.e., 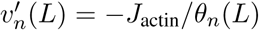. In this example, the actin velocity at the cell front in the fixed frame is 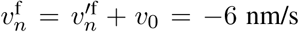. This negative velocity pointing inwards is the actin retrograde flow motion commonly seem during cell migration [31]. Our model also predicts a decrease of the retrograde velocity towards the back of the cell (Fig. 1G,H) as seen in experiments. Because of the boundary condition specified in the model, we do not consider an anterograde actin flow at the back of the cell [6].

### 3.3 Cell migration on 2D substrates relies on actin dynamics and focal adhesion

For cells on 2D substrates the coefficient of hydraulic resistance is negligible (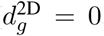, see the SM for more information) and the effective drag from the nucleus on the two phases can be neglected as well (*ζ*_*c*_ = *ζ*_*n*_ = 0). Hence, Eq. 2 is reduced to a simpler form where *v*_0_ is independent of *J*_water_. The model predicts that *v*_0_ scales with the rate of actin polymerization, *J*_actin_*/*Θ*_n_*, and the coefficient of focal adhesions, *η*_st_ (Fig. 2A). In the limit of large *η*_st_ (for example, along the line *η*_st_ = 10^4^ Pa*·*s/*µ*m^2^), the effect of *ξ*_*m*_ and 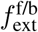 diminishes and *v*_0_ *∼ J*_actin_*/*Θ*_n_*, suggesting that when the strength of focal adhesion is high (no slip), the cell velocity is determined by the rate of actin polymerization. On the other hand, when the force from focal adhesions is abolished (along the line *η*_st_ = 1 Pa*·*s/*µ*m^2^), *v*_0_ does not increase as *J*_actin_ increases. This is consistent with the finding that for actin polymerization to be effective, there must be sufficient focal adhesion friction [19]. Note that the model predicts a monotonic dependence of *v*_0_ on *η*_st_ because we have assumed a linear force-velocity relation for focal adhesions, as opposed to a non-linear relation as seen in experiments [20].

**Figure 2:**
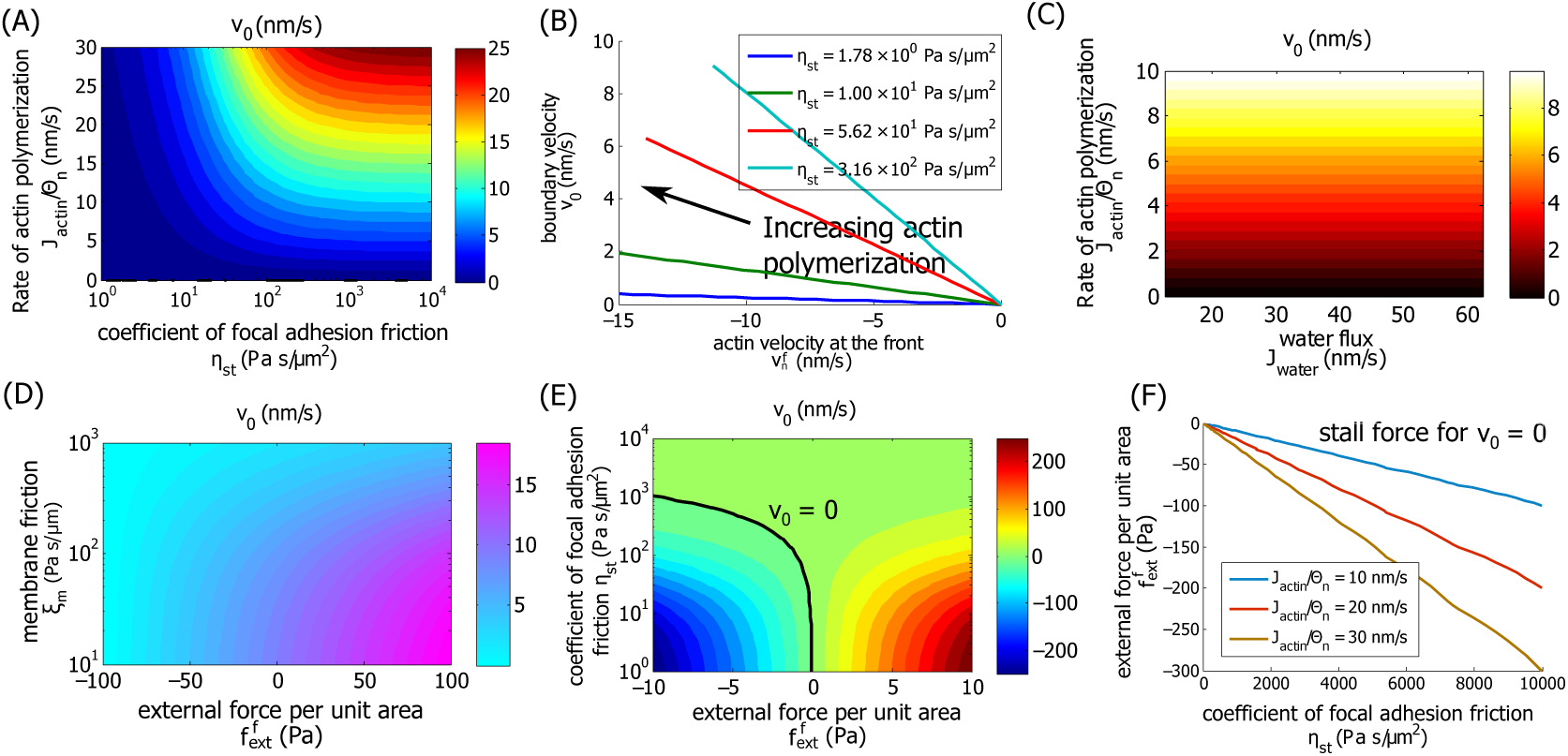
Cell migration on 2D substrates. (A) Contours of the cell boundary velocity, *v*_0_, as functions of actin polymerization rate *J*_actin_ and focal adhesion friction coefficient *η*_st_. *J*_actin_*/*Θ*_n_* is a measure of actin velocity at the cell boundary generated by actin polymerization. In the calculation *J*_actin_ varies and Θ*_n_* remains unchanged. (B) Cell boundary velocity *v*_0_ as a function of 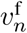 for different coefficient of focal adhesion *η*_st_. (C) Contours of *v*_0_ as *J*_actin_ and *J*_water_ vary. In 2D *J*_water_ does not influence cell boundary velocity. (D) Contours of *v*_0_ as *ξ*_*m*_ and 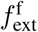 vary. (E) Contours of *v*_0_ as 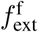 and *η*_st_ vary. (F) Stall force per unit area for different *η*_st_. As expected, the magnitude of the stall force increases with increasing coefficient of focal adhesion friction or the rate of actin polymerization.

Fig. 2B shows the predicted relation between *v*_0_ and actin retrograde velocity as *J*_actin_ varies for different *η*_st_. Higher rate of actin polymerization results in a higher cell velocity and also higher magnitude of actin retrograde flow, since 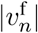 increases faster than *v*_0_. For the same actin velocity *v*_*n*_, the model predicts a higher *v*_0_ for higher *η*_st_.

Although the cell boundary velocity is in general influenced by the water flux through the membrane (Eq. 1), the model predicts that under negligible hydraulic resistance 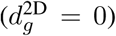 the boundary velocity is independent of *J*_water_. This can also been seen from Eq. 2 where *J*_water_ does not contribute to *v*_0_ when *d*_*g*_ = 0. More discussions are given in a later section.

### 3.4 Influence of external force on cell migration

The cell boundary moves faster if the membrane friction with the environment, *ξ*_*m*_*v*_*m*_, is low, or the combination of the external forces (per unit area), 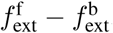, points in the direction of cell migration (Fig. 2D). Contour plots of *v*_0_ as a function of *η*_st_ and 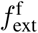 shows a similar story (Fig. 2E). We predict that when the friction force from focal adhesion is small (low *η*_st_), *v*_0_ scales linearly with 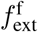; when the force from focal adhesion is large (high *η*_st_), then the external forces has a negligible effect on cell velocity. Therefore, Fig. 2E also implies an external force-cell velocity relation for cell migration. Similar calculation can be obtained for different rate of actin polymerization. We can extract the stall force when *v*_0_ = 0 for each *η*_st_ and *J*_actin_. As expected, the magnitude of the stall force increases as *η*_st_ or *J*_actin_ increases (Fig. 2F). For a 2D cell’s boundary element with 3 *µ*m in width and 200 nm in height, the predicted stall force is on the order of 1 nN, which is of the same order as the stall force for cell boundary lamellipodium [25].

The spatial variation of the cell membrane tension is set partially by the membrane friction with the environment and partially by the external force: larger *ξ*_*m*_ and 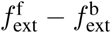 can generate a membrane tension difference up to 10% from the back to the front of the cell (see Fig. S3). In other cases the membrane tension is almost uniform across the cells, and thus the velocity contributed by the membrane trafficking is negligible. This result validates the assumption of *v*_*m*_ = *v*_0_ that is used to approximate *v*_0_ in Eq. 2. The viscosity of each phase or the friction between two phases has little contribution to *v*_0_ (see Fig. S3), showing that for cells on 2D substrates, migration is mostly driven by actin polymerization, focal adhesion, and external forces acting on the cell.

### 3.5 Confined 1D cells migrate faster under higher coefficient of hydraulic resistance

For cells in confined channels (Fig. 3A), the coefficient of hydraulic resistance is 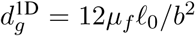, where *£*_0_ is the total length of the channel and *µ*_*f*_ is the extracellular fluid viscosity in the channel (see the SM for more information). A longer channel or a higher extracellular fluid viscosity leads to a higher coefficient of hydraulic resistance. In a typical channel in this work or in [7], for example, *b* = 5 *µ*m, *£*_0_ = 500 *µ*m, and *µ*_*f*_ = 10*^-3^* Pa*·*s, then 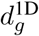 is about 2.5 *×* 10*^-1^* Pa*·*s/*µ*m. In reality, 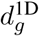 can be much higher than this estimate because the channel walls are not smooth. Fig. 3B shows the model prediction that *v*_0_ increases with 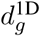 while the other parameters remain the same. The cell velocity saturates at low and high 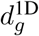. For intermediate 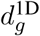, one order of magnitude change of 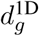, for example from 5 *×* 10^3^ Pa*·*s/*µ*m to 5 *×* 10^4^ Pa*·*s/*µ*m, results in an increase in cell velocity, from *∼*54 *µ*m/hr to *∼*75 *µ*m/hr, by about 40%.

**Figure 3:**
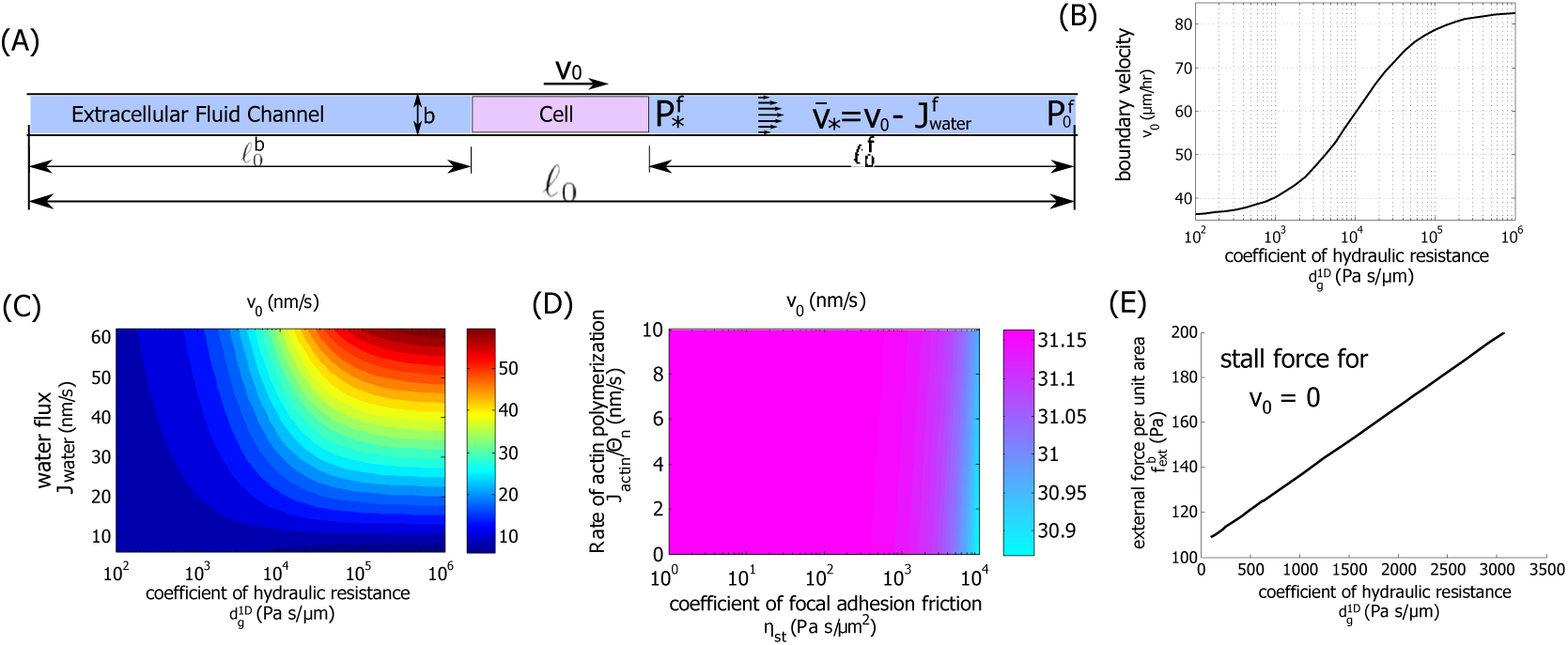
Confined 1D cells migrate faster under higher coefficient of hydraulic resistance. (A) Diagram of the cell and the external fluid flow in a 1D channel. (B) Model prediction of the cell boundary velocity, *v*_0_, as 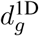 varies. Here 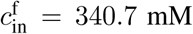 which corresponds to a water flux of *J*_water_ = 82 *µ*m/hr. (C) Velocity of the cell edge, *v*_0_, as *J*_water_ and 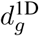 vary. We let vary from 340.6 mM to 341 mM and obtain *J*_water_ accordingly. Cell velocity increases with increasing *J*_water_ and *d*^1D^. (D) Contours of *v*_0_ as *J*_actin_ and *η*_st_ vary. Here 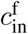. For confined channels with high hydraulic resistance, neither actin polymerization nor focal adhesion friction influence the cell speed significantly. (E) Stall force per unit area increases with 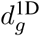.

This prediction, showing that cells move faster in an environment with higher coefficient of hydraulic resistance, is rather counterintuitive. This result can be understood by considering the flow of the extracellular fluid. In Fig. 3B, with constant *J*_water_, *v*_0_ increases as 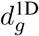 increases, meaning that the cell utilizes more of the water flux in *v*_0_. This means the fluid velocity in the channel, *v*_*\**_ = *v*_0_ *- J*_water_, decreases with 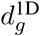, which is expected under higher hydraulic resistance. In addition, the cytosol velocity *v*_*c*_ must be continuous with extracellular fluid velocity and decrease for larger 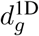. Therefore, the cell boundary velocity increases.

On the other hand, increased velocity of cell migration under higher coefficient of hydraulic resistance, 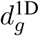, helps to reduce the otherwise high hydraulic resistance, 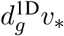: although the coefficient of hydraulic resistance varies from 10^2^ to 10^6^ Ps*·*s/*µ*m, as shown in Fig. 3C, the actual hydraulic resistance experienced by the cell remains in the same order of magnitude due to the increased cell velocity *v*_0_ (see Fig. S4).

### 3.6 Transition between actin-driven and flow-driven cell migrations depends on the external coefficient of hydraulic resistance

Eq. 1 suggests that *v*_0_ increases with *J*_water_; indeed, the model predicts that cell migrates faster under larger water flux. Fig. 3C shows the dependence of *v*_0_ with *J*_water_ and 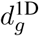. When 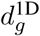 is small 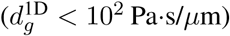, the cell velocity is low and is almost independent of *J*_water_, suggesting that water flux does not contribute to cell migration in this regime. This is the situation for cells on a 2D substrate (also see Fig. 2C). When 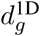 is large 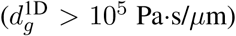, the cell velocity equals to *J*_water_; this is the situation for a cell in a confined space where cell migration is mostly facilitated by water flux, i.e., an osmotic engine model [7]. In this regime, the boundary velocity saturates with 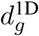.

For high coefficient of hydraulic resistance, for example 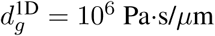, the predicted cell velocity *v*_0_ is independent of *J*_actin_ or *η*_st_ (Fig. 3D). This result, together with Fig. 3C, indicates that under high coefficient of hydraulic resistance, *v*_0_ is dominated by water flux, not the actin network, whereas under low coefficient of hydraulic resistance, *v*_0_ is dominated by the actin network, not water flux. Our model shows that the transition from actin-driven cell migration to water-driven cell migration can result from the physical effects of fluid flow, not from a change in the mechanism of migration signaled by the cell.

Similar to the 2D case where higher external forces are needed to stop cell migration under higher strength of focal adhesion (Fig. 2F), in 1D higher external forces are needed to stop cell migration under higher coefficient of hydraulic resistance (Fig. 3E). The magnitude of the stall force is therefore also a measure of the strength of the ‘driving force’, which is focal adhesion and actin polymerization in 2D and water flux in 1D.

### 3.7 Cell migration in 3D matrices

The model can be extended to understand environmental effects on cell migration in 3D matrices (hydrogel such as collagen) (Fig. 4A). The front of the protrusion interacts with the matrix and the interstitial fluid. As the protrusion extends, it exerts a force on its environment and changes the hydrodynamic pressure distribution in the matrix. The extracellular matrix is a poroelastic material with high porosity. The elastic component, i.e., the collagen fiber network, deforms as the cell protrudes and therefore exerts an external force on the migrating cell. In the case of collagen, the matrix is also dissolved as the cell migrates and secretes metalloproteinases (MMPs) [32].

**Figure 4:**
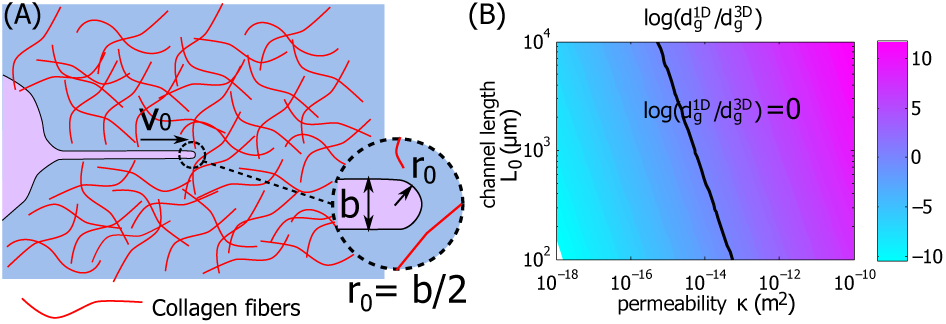
Cell movement in 3D collagen matrix. (A) Diagram of a cell protrusion and the surrounding collagen matrix. Collagen fibers are distributed in the matrix. The tip of the protrusion has a radius *r*_0_ = *b/*2, where *b* is the width of the protrusion. (B) Contour of 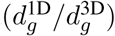 as the 1D channel length *£*_0_ and the permeability of the collagen matrix *κ* vary. The line is where 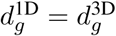.

Using poroelastic theory, we estimate the effective coefficient of hydraulic resistance to be 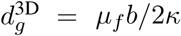 (see the SM for more information), where *µ*_*f*_ is the effective viscosity of the fluid phase in the collagen matrix, *κ* is the permeability of the collagen matrix, and *b* is still the width of the cell protrusion. To differentiate the notation used in 1D, we will add ‘3D’ or ‘1D’ to the corresponding quantities in each geometry, i.e., 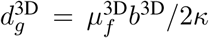 and 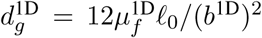. We assume 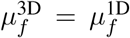. Depending on the values of *b*^3D^*/*2*κ* and 12*£*_0_*/*(*b*^1D^)^2^, the coefficient of hydraulic resistance in 3D can be either larger or smaller than that in 1D. For example, when *κ* = 10*^-16^ m^2^* [33], *b*^3D^ = *b*^1D^ = 3 *µ*m, and *£*_0_ = 10^3^ *µ*m, then 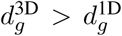. But when *κ* = 10*^-12^ m^2^*, then 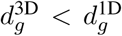. Figure 4B shows the phase diagram of the ratio 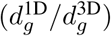 as the 1D channel length *£*_0_ and the permeability of the collagen matrix *κ* vary. The line 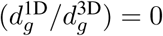 indicates 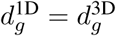.

## 4 Concluding Remarks

Understanding the mechanism of cell migration in different physical environments helps to uncover patho-physiological mechanisms. Using a two-phase modeling framework, we derived a general expression that is applicable to a wide range of physical conditions to compute the cell boundary velocity as a function of actin polymerization rate and water influx rate. The model incorporates known mechanics of important motility components, and includes effects of focal adhesion and membrane friction force, as well as hydraulic resistance from flow outside of the cell. Depending on the environmental properties, the coefficient of hydraulic resistance can be substantial, especially in 1D channels and 3D matrices. The coefficient of hydraulic resistance can influence the relative contribution of water influx to migration speed. In 2D environments, the hydraulic resistance is negligible so that even if the water influx is large, it does not contribute to cell speed as long as the cell maintains constant volume. In 1D confined channels with high coefficient of hydraulic resistance, water influx enhances migration speed. In 3D matrices, the hydraulic resistance depends on the local environment of the cell, and can be as large as the 1D case. This result suggests that in 3D, knocking out cell components responsible for water flux would have major effects in cell motility.

Here we have studied the coefficient of hydraulic resistance over several orders of magnitudes. This large variation range can cover the difference among different types of tissues or oranges and in physiological or pathological conditions. The model prediction suggests that under pathological conditions cells may adopt alternative mechanisms of migration comparing to those migrate under physiological conditions.

Our results express the cell speed for given actin polymerization and water influx rates. However, exactly how these fluxes are controlled is not addressed, and requires additional study. In particular, there likely exists substantial crosstalk between actin polymerization and water flux. Actin polymerization is linked to polarity of the cell [34], and likely influences ion channels and pumps that sets up the osmotic gradient [35]. There are also direct interactions between ion channels and the cytoskeleton [36]. Water and ion fluxes could also reorganize the cytoskeleton. This is especially true for calcium, which is implicated in activating myosin contraction [37]. Therefore, examining the interplay between actin dynamics and water flux would yield new insights.

